# Forensic Identification of Confiscated Helmeted Hornbill (*Rhinoplax vigil*) Casques and Implications for Individual Quantification in Wildlife Crimes

**DOI:** 10.64898/2026.04.02.715475

**Authors:** Ying Shen, Wenzhi Wang, Lianghua Huang, Jing Chen, Kai He

## Abstract

In wildlife forensic practice, species identification and estimation of the Minimum Number of Individuals (MNI) for highly processed specimens have long relied on weight-based conversion methods, which may result in underestimation of the number of individuals involved in a case. Focusing on 41 confiscated casque products of the helmeted hornbill (*Rhinoplax vigil*), including 32 newly added Tianjin samples, this study combines macroscopic morphological examination with mitochondrial DNA barcoding (16S rRNA, COI, and Cytb) to explore a more robust approach for individual quantification. The results demonstrate that the conventional “weight-based” approach overlooks critical biological information contained in anatomical structures and cannot accurately reflect the actual number of individuals involved. Based on this, we propose an anatomy-based criterion centered on the principle of structural uniqueness: specimens retaining biologically unique beak or casque structures should be directly assigned to a single individual, whereas weight-based estimation should only be applied when original anatomical features are entirely absent. In addition, based on morphometric calibration using a larger sample size (n = 40), we propose updating the reference benchmark for estimating the number of individuals in heavily processed solid casque products from 86 g to 65 g. This approach improves the scientific rigor and accuracy of forensic identification and provides reliable technical support for the conviction, sentencing, and law enforcement of wildlife trafficking cases involving helmeted hornbill and other endangered species.

## 1 Introduction

The helmeted hornbill (*Rhinoplax vigil*) belongs to the family Bucerotidae and is one of the largest species within this group. It is primarily distributed in the tropical rainforests of Southeast Asia, including Myanmar, southern Thailand, Malaysia, Indonesia, and Brunei^[1]^. The most distinctive morphological feature of this species is the casque, a helmet-like keratinous structure located on the dorsal side of the upper bill. Unlike the casques of most hornbill species, which are hollow or spongy in structure, the casque of the helmeted hornbill is solid and composed of extremely dense keratin. Its exterior typically exhibits a striking mixture of reddish-brown and yellow. In the illegal wildlife trade, it is commonly referred to as “He Ding Hong”, “hornbill ivory”, or “red ivory”^[2][3]^.

It is precisely this dense texture and uniquely colored solid casque that has rendered the helmeted hornbill a persistent target of illegal hunting and transnational trafficking^[4]^. The casque is frequently carved into high-value luxury items, including snuff bottles, rings, and pendants^[5]^. In the indigenous cultures of Southeast Asia, particularly among Dayak communities in Borneo, the species is revered as a “sacred bird” or a “messenger” linking the human world and spiritual realms. However, in illicit antique markets, such cultural beliefs are often appropriated and misrepresented as “Buddhist sacred objects” with exaggerated claims of properties such as warding off evil, attracting wealth, or even enabling spiritual communication, thereby artificially inflating their market value.Driven by the substantial profits of illegal trade, the helmeted hornbill has experienced severe and unsustainable exploitation. Multiple quantitative assessments underscore the scale of this crisis: between 2010 and 2017, at least 59 trafficking cases were recorded globally, resulting in the seizure of 2,878 casques and related products^[7]^; between 2018 and 2021, a further minimum of 690 individuals and their body parts were confiscated worldwide^[8]^. Reflecting this dramatic decline, the International Union for Conservation of Nature (IUCN) in 2015 elevated the species’ conservation status directly from Near Threatened (NT) to Critically Endangered (CR), skipping intermediate categories^[9]^. Moreover, the species is listed in Appendix I of the Convention on International Trade in Endangered Species of Wild Fauna and Flora (CITES), under which all forms of international commercial trade are strictly prohibited^[10].^

In the forensic practice of wildlife crime investigations, accurate species identification and the estimation of the Minimum Number of Individuals (MNI) constitute critical evidence for conviction and sentencing. However, samples of helmeted hornbill involved in such cases are typically detached from intact individuals and often subjected to varying degrees of physical processing, resulting in the loss or degradation of key morphological characteristics. Under current forensic evaluation guidelines, when confronted with incomplete or heavily processed specimens, the estimation of individual numbers frequently relies solely on sample weight (e.g., specimens weighing less than 100 g may be counted as one-third of an individual). This weight-based approach overlooks essential biological information embedded in morphological structures. Given that helmeted hornbill products undergo substantial material removal during processing, reliance on weight alone is likely to result in a significant underestimation of the number of individuals involved. which may in turn lead to disproportionately lenient sentencing for offenders engaged in poaching and trafficking, thereby failing to achieve an effective deterrent effect of the law.

On the basis of the previous forensic examination of nine suspected helmeted hornbill casque samples, this study further incorporated 32 samples seized in Tianjin, expanding the study material to 41 suspected helmeted hornbill casque products. Species identity was confirmed through mitochondrial DNA (mtDNA) sequencing, and the criteria for determining the number of individuals involved under different processing conditions were systematically evaluated by integrating the morphological integrity of the casque and bill structures. Compared with the previous exploratory study, the present study recalibrated the weight-based reference value for heavily processed products using an expanded sample size and further refined the MNI estimation framework centered on “anatomical structural uniqueness,” with weight-based conversion used only as a supplementary approach. This study aims to provide a more reliable scientific basis for species identification, estimation of the number of individuals involved, and value assessment of helmeted hornbill and other endangered avian products in forensic practice.

## 2 Materials and Methods

### 2.1 Morphological approach for estimating the number of individuals

All submitted samples were subjected to systematic macroscopic morphological examination. Particular attention was paid to the integrity of the casque–bill structure, including the preservation of key features such as the bill tip, bill margins, cutting edges, and the dorsal contour of the casque. Surface coloration characteristics were also recorded, especially the boundary between the reddish-brown region of the casque and the yellowish-white region of the bill. Each sample was weighed using an analytical balance and documented photographically.

This study proposes the anatomical uniqueness of the casque–bill structure as the primary criterion for the determination of the MNI. Given that each helmeted hornbill individual possesses a single solid casque structurally integrated with the upper bill, any processed sample retaining identifiable intact or partial native bill structures, such as a continuous casque-bill structure, identifiable upper bill structure, bill tip, bill margins, or specific keratinous transition zones between the casque and bill, was forensically interpreted as originating from one distinct wild individual, regardless of the residual weight of the processed sample. In contrast, if a sample retained only casque blocks, keratin fragments, beads, plaques, or similar processed materials, and the native bill structure and its original continuity with the casque had been completely lost, making it impossible to determine morphologically whether the sample corresponded to an independent individual, the principle of anatomical structural uniqueness was no longer applicable for direct MNI determination. Such samples were instead classified into the category of weight-assisted estimation.

### 2.2 Sample collection and DNA extraction

A total of 41 processed casque samples suspected to originate from helmeted hornbill were collected in this study, all of which were derived from items seized by Chinese customs authorities during law enforcement operations. The samples were divided into two batches: the first batch consisted of nine samples seized during the earlier stage, whereas the second batch comprised 32 samples subsequently seized in Tianjin. Among the 32 Tianjin samples, 31 retained both casque and bill structures, while only one sample, designated tianjin5, retained only the casque, with the bill completely absent.

Prior to sampling, each sample was assigned a unique identification number, and its morphological characteristics and seizure-related information were recorded. All experimental apparatus were sterilized by autoclaving and ultraviolet irradiation within a dedicated forensic clean bench. The surfaces of the specimens were thoroughly cleaned using 75% medical ethanol and sterile water to remove potential exogenous contamination. Internal keratin debris was then scraped from non-display surfaces of each sample using a sterile scalpel; approximately 500 mg of material was collected per sample and transferred into sterile centrifuge tubes.

DNA was extracted from the samples using the QIAGEN DNeasy Blood and Tissue Kit, and the lysis step was appropriately prolonged for the keratin-rich hard tissues. Following the addition of Proteinase K, samples were incubated at 56 °C for overnight digestion (approximately 12-16 h) until complete lysis was achieved or a homogeneous suspension was obtained. Throughout DNA extraction and subsequent PCR procedures, extraction blanks and PCR negative controls were strictly included to monitor potential cross-contamination.

### 2.3 Gene amplification and sequencing

The extracted DNA was used to amplify mitochondrial 16S rRNA, cytochrome c oxidase subunit I (COI), and cytochrome b (Cytb) gene fragments. The primers used for PCR amplification were as follows:

The 16S rRNA fragment was amplified using primers 16S-F (CGCCTGTTTAYCAAAAACAT) and 16S-R (CCGGTYTGAACTCAGATCAYGT)^[12]^.

Universal primers were employed for COI amplification CO1(TYTCWACWAAYCAYAAAGAYATTGG); CO2 (AYTCAACAAATCATAAAGATATTGG); CO3 (ACYTCYGGRTGACCAAARAAYCA); CO4 (ACYTCRGGRTGACCAAAAAATCA)^[13]^.

The Cytb gene was amplified using primers CYTB-F (CCATCCAACATCTCAGCATGATGAAA) and CYTB-R (CCCTCAGAATGATATTTGTCCTCA)^[14]^.

PCR products were examined by agarose gel electrophoresis, and samples showing a single, clear target band were submitted to a commercial sequencing facility for bidirectional Sanger sequencing. All obtained sequences have been deposited in the GenBank database (accession numbers are provided in **Table 1**).

**Table 1.**
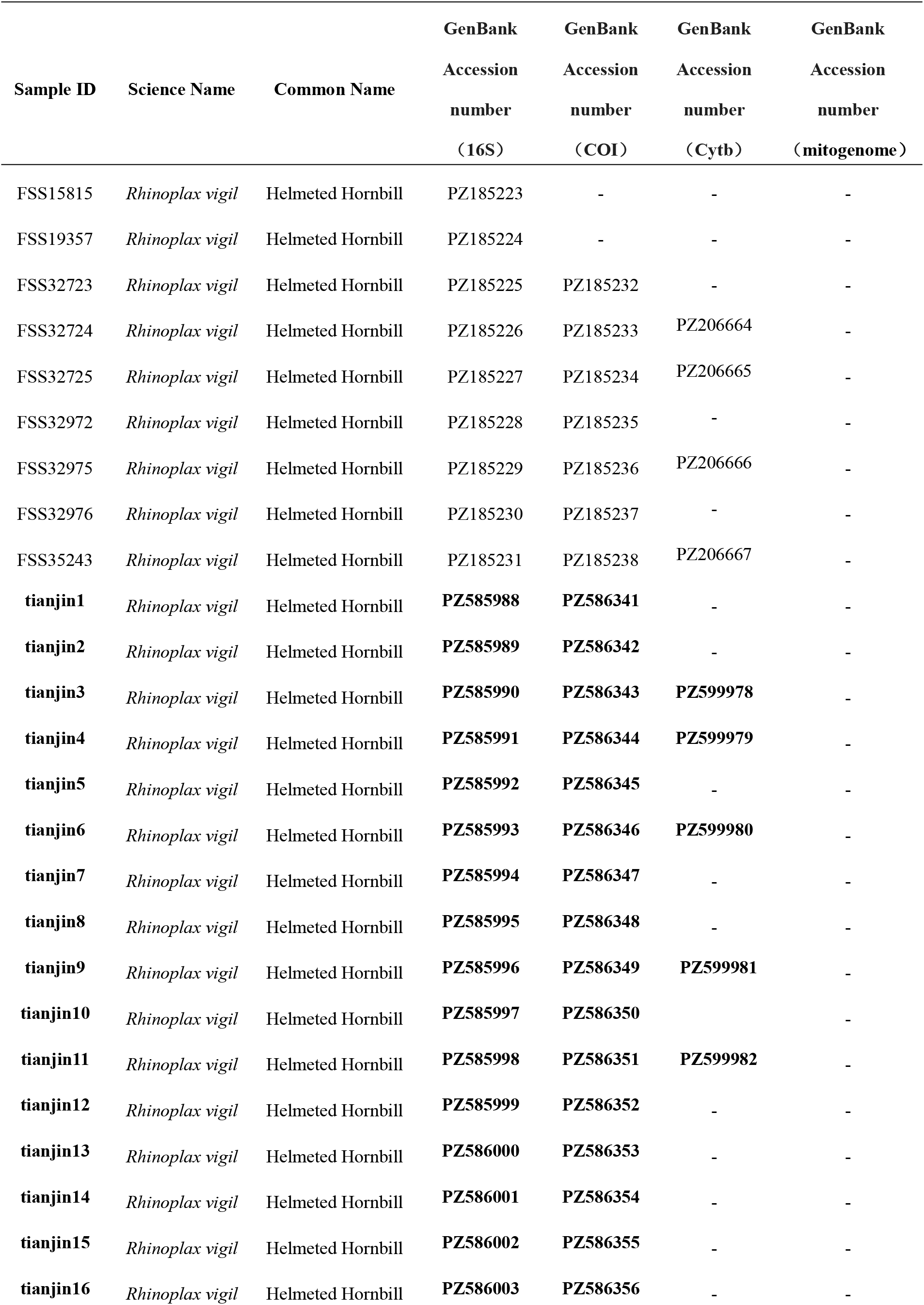
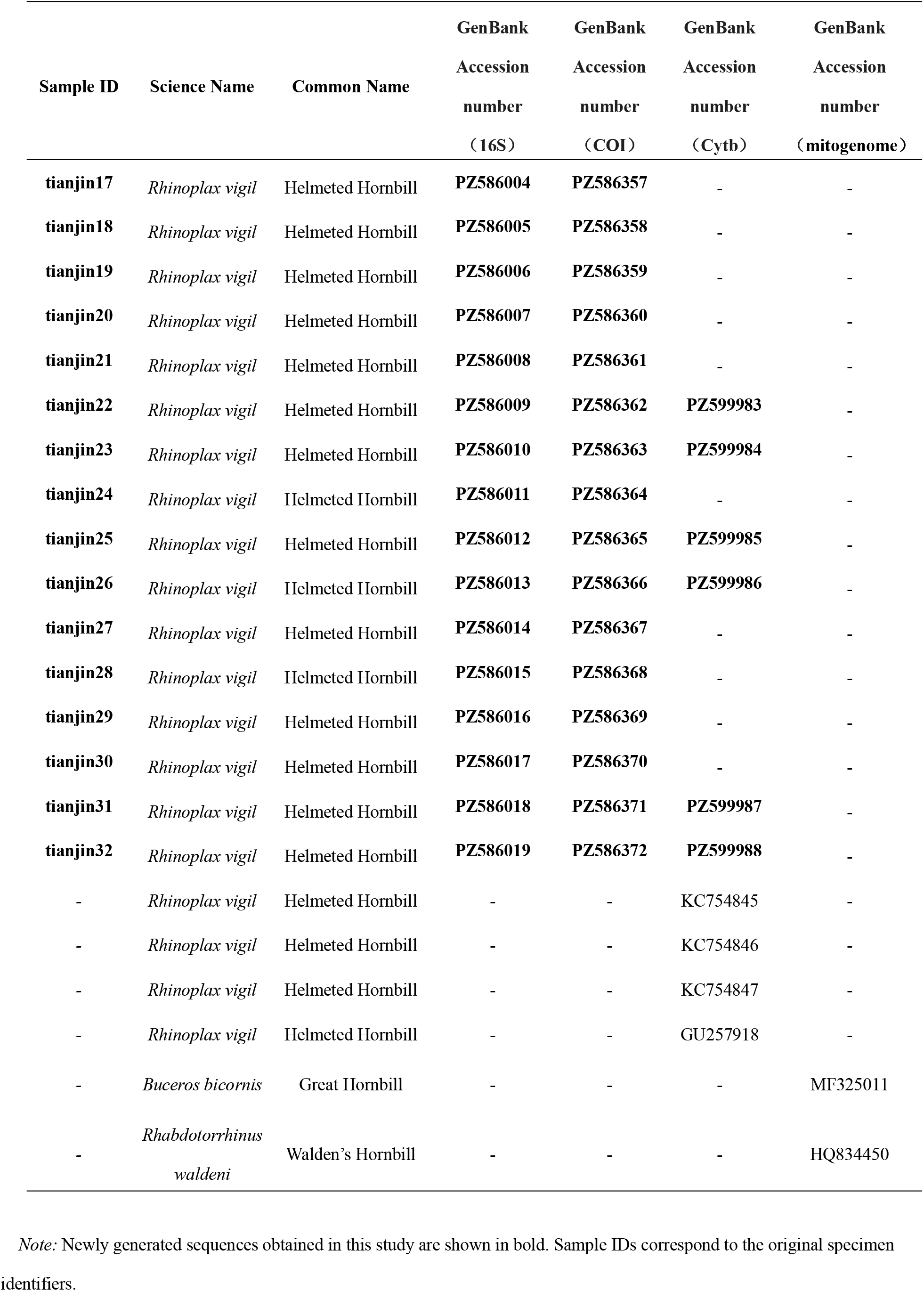
Mitochondrial Gene Sequences (16S rRNA, COI, and Cytb) and Complete Mitochondrial Genomes Used in This Study, With GenBank Accession Numbers and Sample IDs.

### 2.4 Reference dataset construction and phylogenetic analysis

To construct a reliable reference dataset, raw whole-genome resequencing data of the helmeted hornbill were downloaded from the NCBI Sequence Read Archive (SRA; accession no. SRR34218485). De novo assembly of the mitochondrial genome was performed using MitoZ^[15]^, yielding a complete mitochondrial genome sequence, which was subsequently annotated in Geneious Prime v2025.0.2^[16]^. In addition, complete mitochondrial genome sequences of closely related species, including the great hornbill (*Buceros bicornis*) and the Walden’ s hornbill (*Rhabdotorrhinus waldeni*), as well as published Cytb sequences of the helmeted hornbill (accession numbers are provided in Table 1), were retrieved from GenBank and used as references and outgroups for phylogenetic analyses. The obtained sequences were aligned separately for each gene using MAFFT^[17]^. Genetic distances (p-distance) were calculated in MEGA-X^[18]^. Phylogenetic trees were reconstructed using the maximum likelihood (ML) method implemented in IQ-TREE v3.0.1^[19]^, with nodal support assessed by 1,000 bootstrap replicates.

## 3 Results

### 3.1 Morphological characteristics and estimation of the number of individuals

Macroscopic examination revealed that,except for sample tianjin5, the outer surfaces of the casques of the remaining 40 samples exhibited a vivid reddish-brown coloration, smoothly transitioning to yellowish-white toward the bill, and possessed an extremely dense, solid keratin structure (**Fig.1A**), fully consistent with the diagnostic taxonomic characteristics of the helmeted hornbill. Morphological assessment showed that, although these 40 samples had undergone varying degrees of deep processing, including cutting, grinding, and polishing, resulting in substantial loss of original tissue, all samples retained clearly identifiable and non-reproducible anatomical features of the bill. Based on the principle of anatomical uniqueness, forensic interpretation determined that these 40 evidentiary samples originated from at least 40 distinct helmeted hornbill individuals. In contrast, sample tianjin5 retained only the casque portion (**Fig.1B**), and it could not be confirmed whether this sample represented an additional individual. Therefore, it was not included in the morphological MNI calculation and was instead assigned to the category of weight-based estimation.

**Figure 1.**
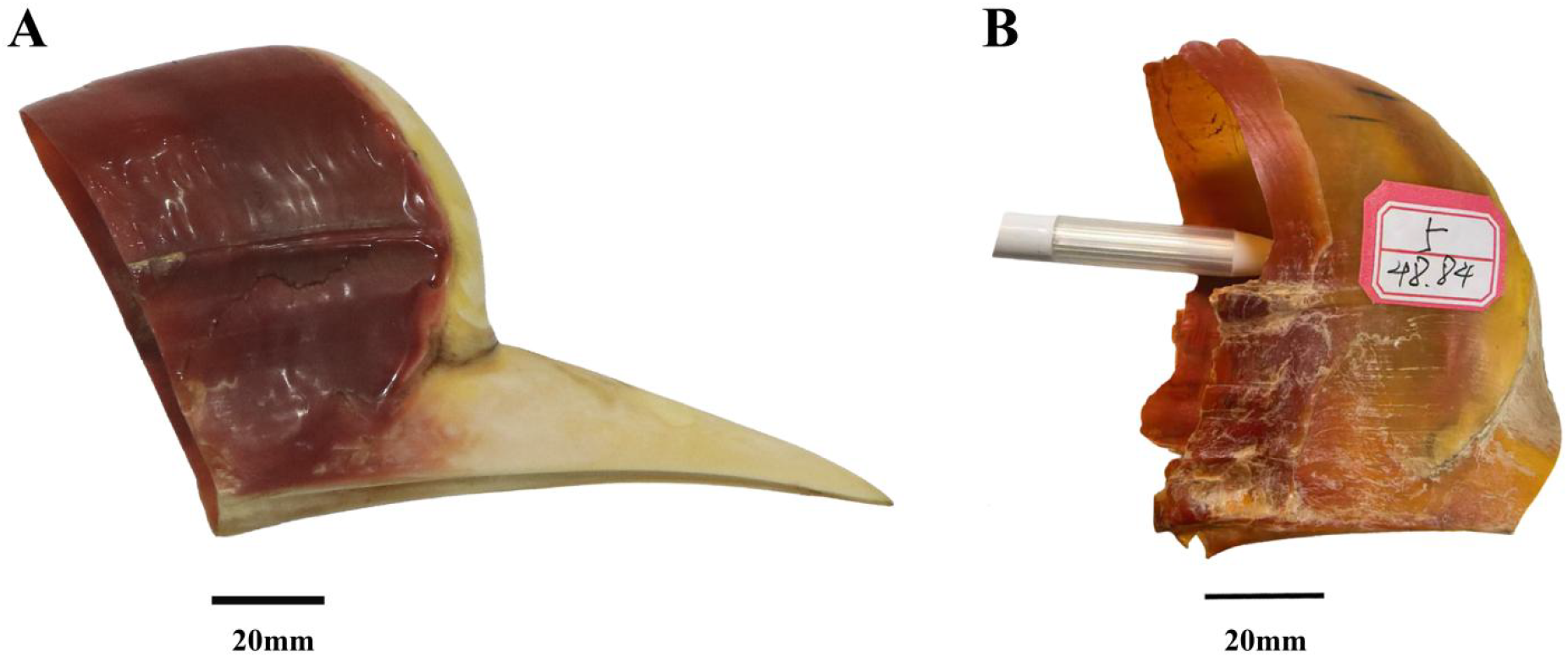
Morphological characteristics of helmeted hornbill (*Rhinoplax vigil*) casque samples. (A)Photograph of a sample retaining the continuous bill–casque structure (No. FSS15815). (B)Photograph of a sample retaining only the casque portion, with the bill absent (No. tianjin5). Scale bars = 20 mm.

Further weight statistics were performed on the 40 samples retaining a complete continuous casque-bill structure. The sample weights ranged from 41.03 to 125.09 g, with a mean of 64.71 ± 16.63 g, a median of 61.47 g, and a 95% confidence interval of 59.40-70.03 g **(Table 2)**. Among them, the nine samples from the first batch ranged from 60.48 to 125.09 g in weight, with a mean of 84.98 ± 18.78 g and a median of 80.40 g, whereas the 31 samples from the second batch ranged from 41.03 to 80.96 g, with a mean of 58.83 ± 10.27 g and a median of 58.24 g. A significant difference was observed between the two batches (Mann - Whitney U test, U = 24.0, p < 0.001), reflecting actual batch differences caused by different seizure sources and processing methods. However, both batches consisted of deeply processed products retaining complete bill – casque structures and could be clearly assigned to single individuals; therefore, they shared a common biological basis for individual determination. To expand the sample size and improve the applicability of the weight-based conversion benchmark to products from different sources, the 40 samples from both batches were pooled for statistical analysis, yielding a mean weight of 64.71 g. Accordingly, the reference value for weight-based conversion of a single deeply processed product was updated from the previous value of approximately 85 g to approximately 65 g. For incomplete samples such as tianjin5, in which the original anatomical structure has been lost, the reference value of 65 g per individual was used for auxiliary weight-based conversion. Thus, 48.84 g ÷ 65 g per individual ≈ 0.75 individual equivalents, which may serve as weight-based evidence when determining the total number of individuals involved in the case.

**Table 2.** Weight statistics of helmeted hornbill (*Rhinoplax vigil*) casque sample.

| Sample ID | Weight (g) | Batch |
| --- | --- | --- |
| FSS15815 | 125.09 | Batch 1 |
| FSS19357 | 94.66 | Batch 1 |
| FSS32723 | 77.18 | Batch 1 |
| FSS32724 | 85.02 | Batch 1 |
| FSS32725 | 80.4 | Batch 1 |
| FSS32972 | 60.48 | Batch 1 |
| FSS32975 | 67.69 | Batch 1 |
| FSS32976 | 78.29 | Batch 1 |
| FSS35243 | 95.97 | Batch 1 |
| tianjin1 | 50.75 | Batch 2 |
| tianjin2 | 58.24 | Batch 2 |
| tianjin3 | 68.89 | Batch 2 |
| tianjin4 | 41.03 | Batch 2 |
| tianjin5 | 48.84 | Batch 2 (Casque only , beak missing) |
| tianjin6 | 64.89 | Batch 2 |
| tianjin7 | 64.48 | Batch 2 |
| tianjin8 | 50.51 | Batch 2 |
| tianjin9 | 54.95 | Batch 2 |
| tianjin10 | 51.84 | Batch 2 |
| tianjin11 | 50.54 | Batch 2 |
| tianjin12 | 49.25 | Batch 2 |
| tianjin13 | 65.71 | Batch 2 |
| tianjin14 | 47.06 | Batch 2 |
| tianjin15 | 62.37 | Batch 2 |
| tianjin16 | 60.56 | Batch 2 |
| tianjin17 | 74.74 | Batch 2 |
| tianjin18 | 71.96 | Batch 2 |
| tianjin19 | 71.94 | Batch 2 |
| tianjin20 | 46.65 | Batch 2 |
| tianjin21 | 47.82 | Batch 2 |
| tianjin22 | 51.36 | Batch 2 |
| tianjin23 | 51.00 | Batch 2 |
| tianjin24 | 73.48 | Batch 2 |
| tianjin25 | 68.28 | Batch 2 |
| tianjin26 | 51.93 | Batch 2 |
| tianjin27 | 51.15 | Batch 2 |
| tianjin28 | 59.90 | Batch 2 |
| tianjin29 | 60.40 | Batch 2 |
| tianjin30 | 71.18 | Batch 2 |
| tianjin31 | 80.96 | Batch 2 |
| tianjin32 | 49.95 | Batch 2 |

### 3.2 Molecular forensic identification

Maximum-likelihood phylogenetic trees were reconstructed based on the 16S rRNA, COI, and Cytb gene sequences, respectively (**Figs. 2A-B**), The three phylogenetic trees consistently showed that the newly added Tianjin batch samples clustered together with the previous first-batch samples and reference sequences of the helmeted hornbill (SRR34218485 and published GenBank sequences), forming a well-supported monophyletic clade with high nodal support values (16S rRNA: 98%; COI: 97%; Cytb: 99%). Meanwhile, all samples were clearly separated from the closely related species *Buceros bicornis* and *Rhabdotorrhinus waldeni*.

**Figure 2.**
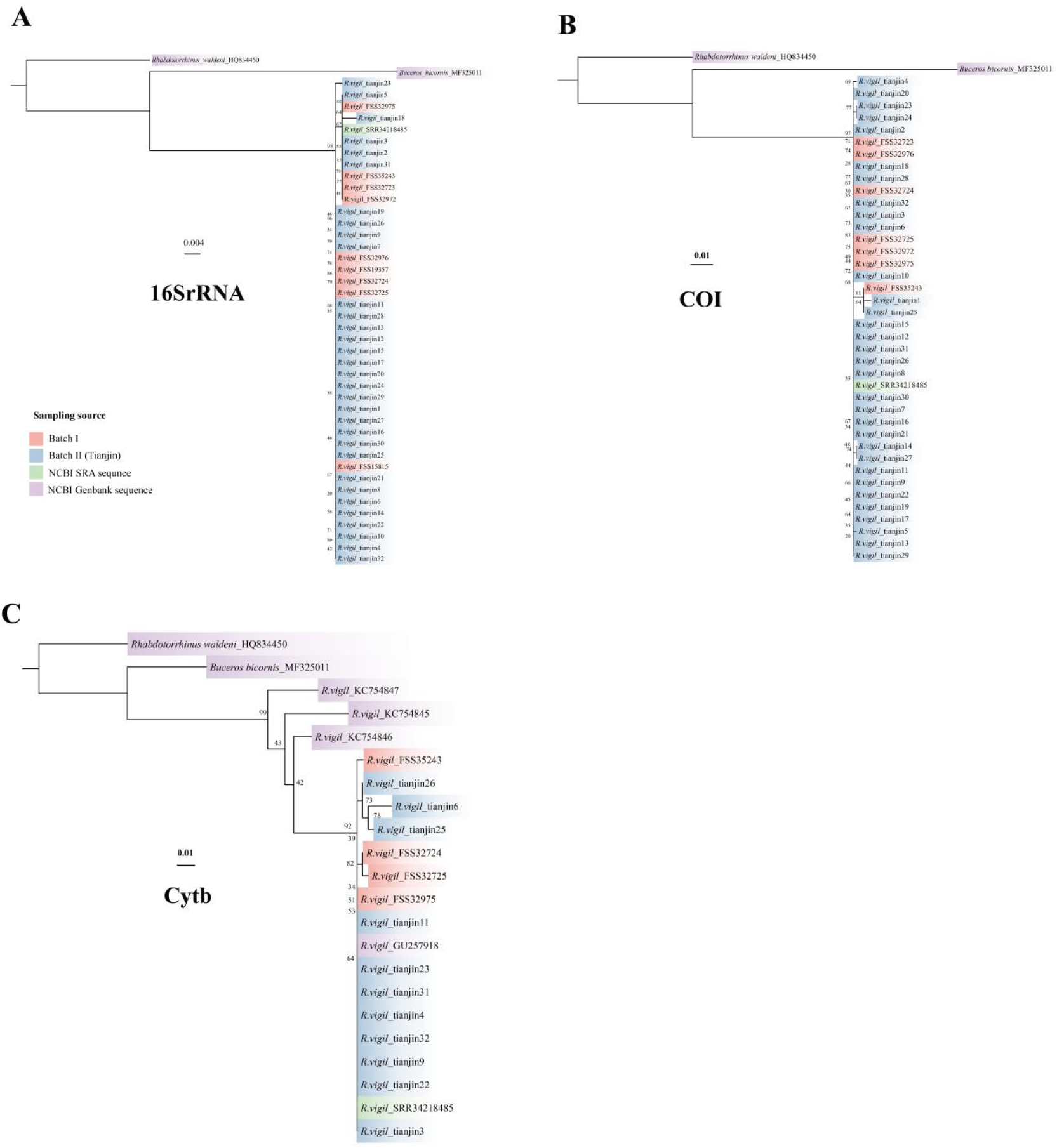
Maximum-likelihood phylogenetic trees of helmeted hornbill (*Rhinoplax vigil*) samples constructed based on mitochondrial genes. (A) 16S rRNA; (B) COI; (C) Cytb. Different colors indicate sample sources: first-batch samples, Tianjin-batch samples, NCBI SRA reference sequences, and GenBank reference sequences. Numbers at the nodes indicate bootstrap support values, and scale bars represent the number of substitutions per site.

Genetic distance analysis revealed extremely low intraspecific divergence between the samples and helmeted hornbill reference sequences (SRR34218485) (16S rRNA: 0.156%; COI: 0.518%; Cytb: 1.855%). When compared with some GenBank reference sequences, however, the Cytb gene showed a certain degree of variation, with pairwise genetic distances reaching up to 10.117% in a few cases. Although this value exceeds the commonly observed range of intraspecific variation, it remains markedly lower than the interspecific divergence between *Rhinoplax vigil* and its closely related species, including *Buceros bicornis* and *Rhabdotorrhinus waldeni* (12.004% and 14.005%, respectively),, and does not cross the evident barcoding gap. Taken together, the molecular evidence unequivocally supports that all 41 evidentiary samples originated from the helmeted hornbill .

## 4 Discussion

By integrating morphological examination with multi-locus mitochondrial DNA barcoding, this study reliably identified the species and estimated the number of individuals represented by highly processed smuggled helmeted hornbill casque products. Compared with the previous exploratory study based on nine samples, the present study further incorporated 32 newly added Tianjin-batch samples, expanding the study material to 41 evidentiary products. Based on this enlarged sample size, the weight-based conversion reference value for heavily processed products was re-evaluated. This study provides important methodological guidance for the field of wildlife forensics, particularly in the species identification of processed products derived from endangered taxa and in the refinement of criteria for estimating the MNI.

### 4.1 Molecular forensic identification strategy for processed helmeted hornbill products

The casque of the helmeted hornbill consists of highly dense keratin and is frequently carved into artifacts such as bracelets and pendants in illegal trade, resulting in the loss of original morphological features. Under such conditions, conventional morphology-based identification becomes challenging. The present study demonstrates that, despite intensive physical processing (e.g., cutting, grinding, and polishing), sufficient DNA can still be recovered from the keratin matrix to enable PCR amplification.

Although previous studies have reported that the phenol-chloroform-isoamyl alcohol (PCIA)^[20]^ method may yield higher DNA recovery from keratinized tissues compared with silica membrane-based commercial kits (e.g., Qiagen). Our results demonstrate that, after rigorous surface decontamination, an optimized protocol using a Qiagen silica column-based kit can consistently provide DNA suitable for short-fragment PCR amplification.

In addition, the helmeted hornbill belongs to a monotypic genus (*Rhinoplax*), lacking closely related sister taxa that could confound species-level identification. Phylogenetic analyses showed that the newly added 32 Tianjin samples clustered tightly with the nine first-batch samples and helmeted hornbill reference sequences, forming the same highly supported clade. Phylogenetic analyses also showed that all three mitochondrial markers (16S rRNA, COI, and Cytb) exhibited clear interspecific divergence from closely related hornbill species in the same family, including *Buceros bicornis* and *Rhabdotorrhinus waldeni*, forming well-resolved genetic boundaries. These results suggest that each of these loci can serve as a reliable molecular marker for the forensic identification of helmeted hornbill^[20][21]^[22][23].

### 4.2 Extremely Low Intraspecific Genetic Diversity and Conservation Warning

In this study, the genetic distances between the samples and reference sequences were extremely low (e.g., 16SrRNA p-distance of only 0.156%). The newly added Tianjin-batch samples and the first-batch samples clustered together within the helmeted hornbill clade in the phylogenetic trees and did not form independent branches corresponding to their seizure batches, suggesting high sequence similarity among samples from different batches in the mitochondrial fragments analyzed. These results are highly consistent with recent genetic studies of helmeted hornbill populations. The elevated genetic homogeneity not only reflects the species’ naturally low population density but also serves as a stark warning of recent population bottleneck effects caused by intensive, large-scale poaching (e.g., approximately 500 individuals reportedly killed per month in West Kalimantan, Indonesia, in 2013^[24]^). The persistent “bleeding” of wild populations by highly organized criminal networks is rapidly eroding the species’ genetic potential to adapt to environmental changes.

### 4.3 “Morphology-First” Approach: Reconstructing Forensic Criteria for Estimating the Number of Individuals

The core focus of this study is how to scientifically determine the MNI in forensic cases. Literature reports^[2][24]^[25] indicate that the solid casque of an adult helmeted hornbill, together with the skull, accounts for only about 10-13% of the body weight (i.e., approximately 270-400 g). During the smuggling process, the removal of the skull, internal cancellous bone, and peripheral fragments of the red-yellow keratin layers results in a precipitous reduction in the weight of the products. Current forensic guidelines, lacking consideration of the species-specific characteristics, mechanically rely on “weight-based conversion rules” (e.g., <100 g counted as 1/3 individual), which contain severe logical flaws. In this case, the 9 first-batch helmeted hornbill samples retained complete upper bill anatomical structures, by virtue of their intrinsic physical properties, provide irrefutable evidence that they originated from nine adult individuals that were killed; however, their mean weight is only 84.98 g. According to the existing weight-based conversion rules, these 9 critically endangered individuals that were brutally killed would legally be reduced to only three individuals in official documentation. This substantial underestimation of individual numbers can lead to downgraded convictions and reduced fines, indirectly facilitating wildlife trafficking.

It is noteworthy that, among the newly added Tianjin batch, 31 samples retained complete continuous casque-bill structures, with weights ranging from 41.03 to 80.96 g and a mean weight of 58.83 g. This differed significantly from the first-batch samples, which ranged from 60.48 to 125.09 g, with a mean weight of 84.98 g, reflecting actual batch differences caused by different seizure sources and processing methods. However, the two batches were completely consistent in the biological fact that each sample corresponded to a single individual, and therefore shared a common basis for individual determination. In this study, the two batches comprising 40 samples retaining complete casque-bill structures were pooled for statistical analysis, yielding a mean weight of 64.71 g and a 95% confidence interval of 59.40 - 70.03 g. Accordingly, the weight-based conversion benchmark for deeply processed products was updated from the previous value of approximately 85 g to approximately 65 g.

Therefore, this study strongly advocates for the following revisions to the current forensic standards:

1. Principle of Anatomical Uniqueness: If a seized sample retains biologically unique anatomical structures (e.g., preserved connection between upper and lower bills, specific casque protuberances), it should be directly recognized as one independent individual regardless of its post-processing weight.
2. Reference Weight for Deeply Processed Products: Only when the sample has been thoroughly cut into beads, plaques, or other fragments, resulting in the complete loss of original anatomical structures, should weight-based conversion be applied. Based on the updated statistical results from 40 helmeted hornbill samples, the mean weight of deeply processed casques retaining basic morphology was 64.71 g. We recommend that, in legal practice, the MNI reference weight for a single deeply processed solid product be updated to 65 g (rather than the full 300 g or the previously provisional value of 85 g). For sample tianjin5, weight-based conversion may be applied using this benchmark; its weight of 48.84 g corresponds to approximately 0.75 individual equivalents. This result may serve as weight-based evidence in the comprehensive determination of the total number of individuals involved and should be listed separately from the 40 individuals confirmed by morphological evidence. This approach accounts for material loss during physical processing and maximizes the accuracy of reflecting the actual scale of poaching committed by offenders.

In conclusion, in the forensic identification of the critically endangered helmeted hornbill, an integrated approach combining morphological integrity assessment with multi-locus molecular tracing represents best practice. Establishing criteria for estimating the number of individuals based on a “morphological anatomical evidence-first” principle, supplemented by post-processing weight-based thresholds, can not only significantly improve the scientific robustness of forensic conclusions but also provide strong technical support, from a judicial perspective, for effectively prosecuting transnational wildlife trafficking networks and curbing illegal demand for this species.

